# *Pseudomonas* synergizes with fluconazole against *Candida* during treatment of polymicrobial infection

**DOI:** 10.1101/2021.11.15.468768

**Authors:** Siham Hattab, Anna-Maria Dagher, Robert T. Wheeler

## Abstract

Polymicrobial infections are challenging to treat because we don’t fully understand how pathogens interact during infection and how these interactions affect drug efficacy. *Candida albicans* and *Pseudomonas aeruginosa* are opportunistic pathogens that can be found in similar sites of infection such as in burn wounds and most importantly in the lungs of CF and mechanically ventilated patients. *C. albicans* is particularly difficult to treat because of the paucity of antifungal agents, some of which lack fungicidal activity. In this study, we investigated the efficacy of anti-fungal treatment during *C. albicans*-*P. aeruginosa* co-culture *in vitro* and co-infection in the mucosal zebrafish infection model analogous to the lung. We find that *P. aeruginosa* enhances the activity of fluconazole (FLC), an anti-fungal drug that is fungistatic *in vitro*, to promote both clearance of *C. albicans* during co-infection *in vivo* and fungal killing *in vitro*. This synergy between FLC treatment andbacterial antagonism is partly due to iron piracy, as it is reduced upon iron supplementation and knockout of bacterial siderophores. Our work demonstrates that FLC has enhanced activity in clinically relevant contexts and highlights the need to understand antimicrobial effectiveness in the complex environment of the host with its associated microbial communities.

## Introduction

Opportunistic microbes co-inhabit host niches, leading to difficult-to-treat co-infections of immunocompromised individuals. However, we know little about how host niche and microbe-microbe interactions affect antimicrobial sensitivity. *Candida albicans* and *Pseudomonas aeruginosa* are two of the most prolific opportunistic pathogens in the developed world, inhabit the same niches and are associated with polymicrobial infections (1–3). *Candida* is the 4^th^ most common nosocomial pathogen and *Pseudomonas* is also associated with significant mono-microbial disease (4–6). *Candida*-*Pseudomonas* co-infections are associated with exacerbated disease, but it is not clear if co-infection should be treated with the same antimicrobials as mono-infection (7–10).

*C. albicans* and *P. aeruginosa* co-colonize numerous sites on the human body, including the gut, lungs, burn wounds, genitourinary tract, but most importantly they can be co-isolated in the lungs of cystic fibrosis (CF) patients (11, 12). CF is a genetic disease characterized by poor mucus clearance in the respiratory tract that leads to persistent infections and polymicrobial biofilms. *P. aeruginosa* infects around 70% of CF patients by the age of 30, and *C. albicans* is isolated in 75% of CF patients (13). Simultaneous colonization by these two pathogens has been linked to more severe clinical outcomes, due to accelerated decline in lung function and worsening of disease progression (7–10). However, the mechanism(s) underlying the postulated enhanced virulence are unknown, so it is difficult to determine if and how this interkingdom dialog regulates pathogenesis and therapy.

Co-infection of *Candida* with diverse bacteria leads to enhanced virulence (11, 22). *C. albicans* and *P. aeruginosa* interact through physical association, secreted factors and signaling molecules that can modulate important virulence factors in both pathogens. *In vitro* studies suggest that antagonistic interactions take place between *P. aeruginosa* and *C. albicans* through phenazines, ethanol and quorum sensing molecules (23–27). Diverse *in vivo* studies of *Candida-Pseudomonas* co-infection have shown either enhanced or decreased virulence (12, 23, 25–28). These *in vivo* studies suggest that a sophisticated understanding of the consequences of co-infection should account for multiple factors such as host environment, nutrient availability and host immune response that might shape these interactions.

*C. albicans* and *P. aeruginosa* have diverse strategies to sense host-relevant cues and adapt their cellular responses based on nutrient availability in the host (14–17). Micronutrient acquisition is a crucial aspect of virulence for most pathogens, including *Candida* and *Pseudomonas* (14, 18). Niche-specific levels of iron even lead to differential dependence on iron sensing and response machinery for *Candida*, with iron-rich environments requiring detoxification and iron-poor environments requiring enhanced acquisition (14, 19). Interestingly, iron starvation has been linked to altered antimicrobial susceptibility (20, 21). Understanding different pathways controlling these adaptation strategies will reveal new opportunities for novel therapeutic targets.

Previous studies have predominantly focused on physical and molecular interactions between *C. albicans* and *P. aeruginosa* and their effect on growth, morphology and virulence, but little is known about effects of cohabitation with antimicrobial treatment. While mixed biofilms enhance antibacterial effects, it is relatively unexplored how these fungal-bacterial interactions affect antifungal drug efficacy during infection (29–31). Fluconazole (FLC) is highly effective and widely used in clinical settings to treat and prevent fungal infections, but paradoxically acts as a fungistatic drug *in vitro* (32). FLC tolerance is high among some clinical isolates and is associated with empirical treatment failure and worse outcomes (33), suggesting that reducing tolerance with adjuvant therapy may boost treatment success. Tolerance is frequently measured as trailing growth and manifests as slow *in vitro* growth of *C. albicans* in the presence of FLC at concentrations above the MIC. Fungicidal activity can be achieved *in vitro* with the addition of drugs such as HMG-CoA reductase inhibitors, calcineurin inhibitors, phenazines or iron chelators (20-21, 33-39). Since microbes naturally produce these types of inhibitors, this raises the possibility that co-colonization or co-infection can produce conditions that enhance FLC activity.

To investigate if *C. albicans*-*P. aeruginosa* interactions affect FLC efficacy, we studied its activity *in vitro* and in the zebrafish infection model. Zebrafish is a powerful model organism that offers the advantage of examining infection outcomes *in vivo* while monitoring host and pathogen physiology through high resolution imaging (40–41). The swimbladder is similar to the human lung, in that they are both air-filled, have a single layer epithelial lining that produces surfactant, and they share similar gene expression patterns (42–46). These similarities make the swimbladder infection model a useful tool to study mucosal infections (23, 47–51).

Previously, we found that *P. aeruginosa* and *C. albicans* are synergistically virulent in the swimbladder model, with enhanced invasive *C. albicans* growth and increased fish mortality (23). In this work, we investigated if FLC efficacy is modulated by *P. aeruginosa* during co-culture and co-infection. Surprisingly, we observed that the combination of *P. aeruginosa* and FLC is synergistic against *C. albicans*, making the drug fungicidal and increasing its efficacy by over 3-logs. This striking effect was seen both *in vitro* and *in vivo.* Interestingly, iron supplementation led to a partial reversal of this synergy *in vitro* and *in vivo*. Taken together, these results suggest that the presence of co-colonizing or co-infecting microbes can substantially affect drug susceptibility in the vertebrate host.

## Results

### Fluconazole is synergistic with *P. aeruginosa* against *C. albicans in vitro*

*C. albicans* and *P. aeruginosa* are common opportunistic pathogens that are found in co-infections at multiple body sites, especially in the lungs of cystic fibrosis patients. We understand little about how co-infection affects virulence or whether treatment should be customized when both bacterium and fungus are co-isolated (13, 31). To determine if interactions between these microbes affect antimicrobial sensitivity, we performed co-culture experiments to test if *P. aeruginosa* affects the antifungal action of fluconazole (FLC) against *C. albicans*. While FLC is fungistatic *in vitro* against *C. albicans*, in *C. albicans* - *P. aeruginosa* co-culture it had potent fungicidal activity. FLC alone slowed growth of *C. albicans* while *P. aeruginosa* alone showed little to no effect on *C. albicans* growth, however the combination led to killing of greater than 1000x from the initial fungal inoculum (Fig. 1 A). This loss of fungal viability is a hallmark for loss of FLC tolerance. Several FLC hyper-resistant *C. albicans* clinical isolates are also susceptible to this FLC*-P. aeruginosa* combination when supra-MIC concentrations of FLC are used (Fig. S1). Synergy was also seen for susceptible and resistant clinical isolates of *C. glabrata*, which is evolutionarily distinct from *C. albicans* and has intrinsic fluconazole resistance (Fig. S1). Fungicidal synergy was not observed with heat-killed bacteria (Fig. S2). The enhancement of FLC activity was reproducible in other media (YPD + serum) and at different temperatures (30° C, 37° C) (Fig. S3). Although these results argue for a robust bacteria-drug synergy, no *in vitro* conditions can truly substitute for the dynamic immune and nutritional environment found during infection.

**Fig. 1:**
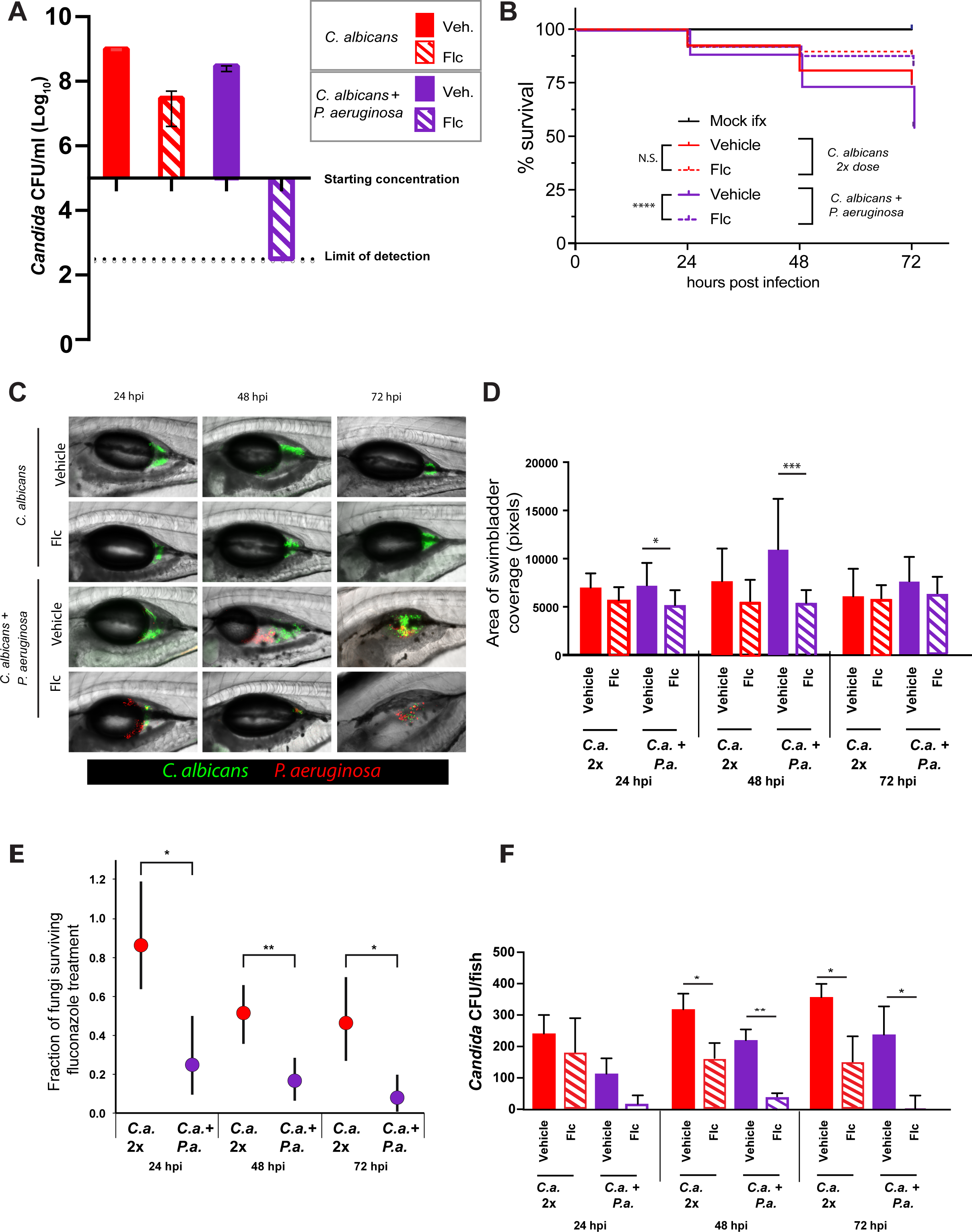
Fluconazole is synergistic with *P. aeruginosa* against *C. albicans*. **A)** *C. albicans + P. aeruginosa* + FLC shows a fungicidal effect after co-culture. *P. aeruginosa* and *C. albicans* were inoculated at 2x10^5^/ml and FLC was added at 12.5 μg/ml. Drops (3 μl) of serial 10x dilutions of co-cultures were plated on YPD containing antibiotics. Representative of >20 independent experiments. **B-D)** FLC is more effective against co-infection. **B)** Fish were infected in the swimbladder with either 50-100 *C. albicans* (mono-infection) or with 25-50 *C. albicans* and 25-50 *P. aeruginosa*, screened for fungal inoculum, then reared in water with or without 100 μg/ml FLC. Data pooled from 13 independent experiments. **C)** Representative images of swimbladder infected with *C. albicans* or *C. albicans* + *P. aeruginosa* with or without FLC (100 μg/ml). Scale bars = 100 μm. Dotted white lines mark the boundary of the swimbladder. **D)** Fraction of fungi surviving FLC treatment. Results are from 5 independent experiments. Monte-Carlo analysis was used to compare groups. **E)** *C. albicans* burden was measured by analysis of confocal z-stacks. Data from 13 independent experiments. Graphs show medians and 95% confidence intervals. Data from 13 independent experiments. **F)** *C. albicans* burden calculated by CFUs. Data from 5 independent experiments. (p >0.05 NS; < 0.05 *; <0.01 **; <0.001 ***; <0.0001 ****)

### *P. aeruginosa* enhances fluconazole activity against *C. albicans* during swimbladder infection in zebrafish

To further examine *C. albicans-P. aeruginosa* interactions in the presence of FLC *in vivo*, we leveraged our zebrafish swimbladder co-infection model. This mucosal co-infection mimics conditions similar to human lungs and leads to synergistic virulence through enhanced fungal invasiveness (23, 50). To induce similar levels of mortality in mono- and co-infection, larvae were either mono-infected with a double dose of *C. albicans* (50-100 cells/fish) or co-infected with *C. albicans* (25-50 cells/fish) plus *P. aeruginosa* (50 cells/fish). We found that FLC treatment significantly reduced mortality in co-infection, although there was only a trend toward reduced mortality in the mono-infected group (Fig 1B). This difference is reflected in different hazard ratios for FLC treatment in monoinfection (0.446, 95% CI 0.257-0.775) and co-infection (0.3255, 95% CI 0.243-0.437). This indicates that FLC is more effective in treating *C. albicans-P. aeruginosa* co-infection than fungal mono-infection, suggesting that there is also bacterial-drug synergy *in vivo*.

The enhanced survival of FLC-treated co-infected fish could be due to effects on the fungus, the bacteria and/or the host. We found that FLC does not affect zebrafish health (Fig. S4) or *P. aeruginosa* growth *in vitro* (Fig. S5). To test if decreased mortality is due to a decrease in *C. albicans* burden, fish were imaged by confocal microscopy at 24, 48 and 72 hpi, and we found fewer fluorescent *C. albicans* cells when co-infections were treated with FLC (Fig. 1C). This burden was quantified by counting fluorescent *C. albicans* pixels in the swimbladder. By this measure, FLC caused no significant decrease in *C. albicans* burden in mono-infected fish, but it caused a significant reduction in co-infected fish at 24 hpi and 48 hpi (Fig. 1D). Additionally, we homogenized fish and measured the number of viable *C. albicans* CFU per fish. The fraction of fungi surviving FLC treatment was strikingly higher during mono-infection compared to the co-infection, while there were almost no viable fungi in the co-infected fish treated with FLC (Fig. 1E & 1F). This CFU data is particularly robust, as FLC inhibits hyphal formation (52–54) and the process of homogenization biases against fungal hyphae, due to their strong inter-hyphal adherence and connections which tend to err on the side of undercounting. Together, these data suggest that the combination of FLC and *P. aeruginosa* have a fungicidal effect against *C. albicans* both *in vitro* and *in vivo*.

### Fluconazole - *P. aeruginosa* synergy is associated with iron limitation

Several molecular interactions between *C. albicans* and *P. aeruginosa* play roles *in vitro* and during infection, including quorum sensing, phenazine toxins, fungal morphogenesis and iron starvation (12). Iron is an important micronutrient for both *C. albicans* and *P. aeruginosa*, and iron chelation leads to enhanced FLC activity against *C. albicans* (20, 38, 55). To determine if *P. aeruginosa* enhances FLC activity against *C. albicans* by outcompeting for iron, we supplemented co-cultures of *C. albicans* and *P. aeruginosa* and FLC with 1 mM FeCl_3_ *in vitro*. We found that iron supplementation limits but does not eliminate the synergistic fungicidal activity of the *P. aeruginosa*-FLC combination (Fig 2A). Similarly, bacteria lacking the two major siderophores pyoverdine and pyochelin had a slightly reduced ability to synergize with FLC, although the rescue was not as strong as with iron supplementation and was not affected by additional iron supplementation (Fig. 2B). The clear loss in fungal viability even upon iron supplementation is consistent with an inability for iron to restore FLC tolerance. Intriguingly, at high concentrations phenazines have been shown to have a synergistic effect with azoles against *C. albicans in vitro* (21). However, both Δ*lasR* and Δ*phz* bacterial mutants had undiminished synergy with FLC against *C. albicans* (Fig. S6). Filamentous fungal growth also does not appear to play a role, as this synergy occurs both in YPD, with >99% yeast, and in RPMI, with >50% hyphae and pseudohyphae (Fig. S3). These results indicate that *P. aeruginosa* synergizes with FLC *in vitro* in part by out-competing *C. albicans* for iron, but this synergy is largely independent of previously identified mediators of *Candida*-*Pseudomonas* dialog (quorum sensing, phenazine toxin production and fungal filamentous growth).

**Figure 2:**
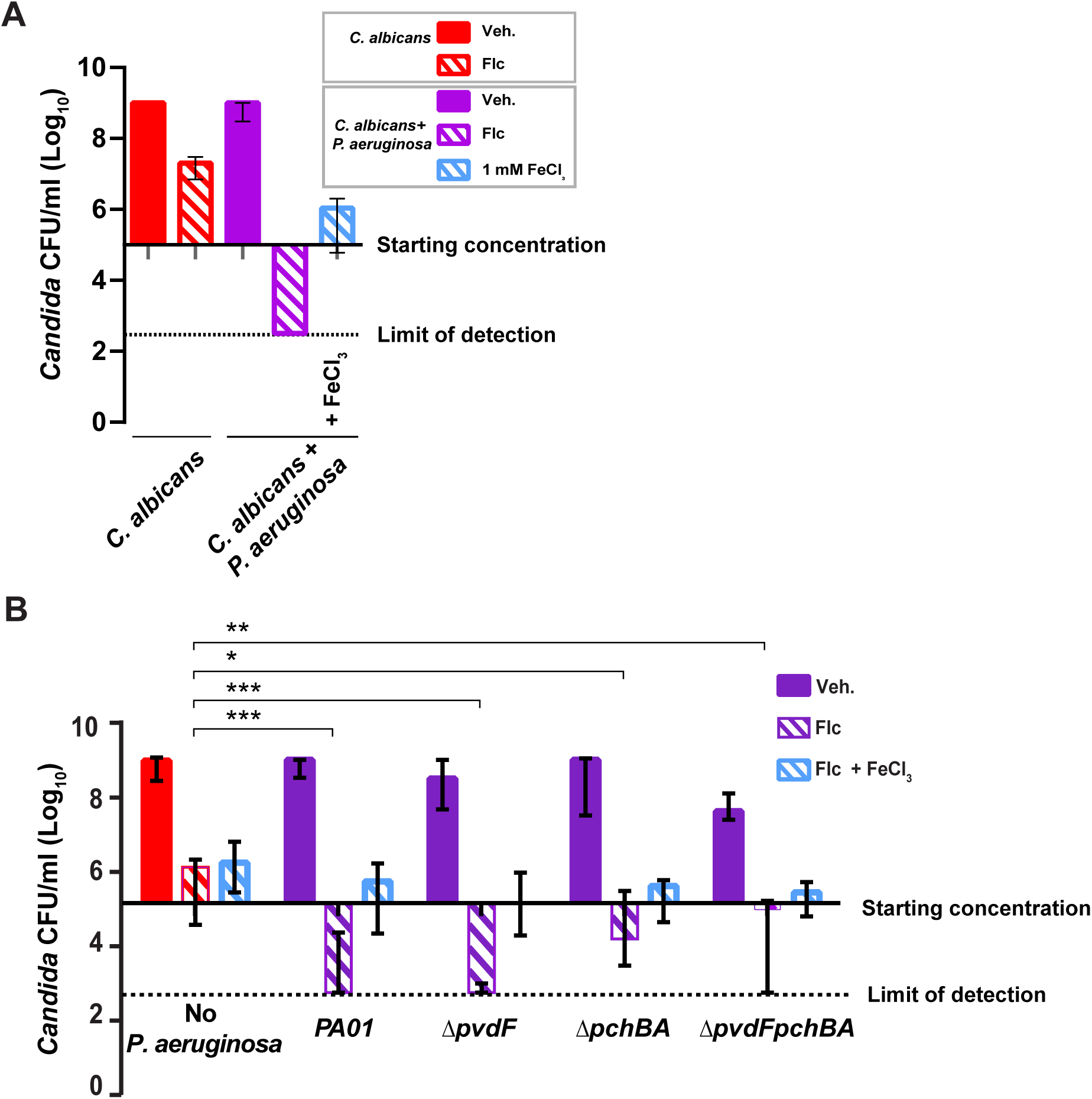
Iron supplementation partially reverses fungicidal effect *in vitro*. **A)** FeCl_3_ supplementation reverses *P. aeruginosa*-FLC synergy *in vitro*. Co-cultures were done with or without FLC treatment (12.5 μg/ml) and/or FeCl_3_ (1 mM). Data from 3 independent experiments. **B)** *C. albicans* growth after 48 h co-cultures with *P. aeruginosa* WT or siderophore mutants: Δ*pvdF, ΔpchB, ΔpvdFpchBA*. Bar graph represents *C. albicans* growth in log_10_ CFU/ml. Data is representative of 4 independent experiments and medians with interquartile ranges from three independent experiments are shown. (p >0.05 NS; < 0.05 *; <0.01 **; <0.001 ***; <0.0001 ****)

### *P. aeruginosa* supernatant in combination with FLC has a partial activity against *C. albicans*

The implication of iron starvation in the interaction between *C. albicans* and *P. aeruginosa* suggested that secreted molecules might drive synergy with FLC. *P. aeruginosa* secretes a large number of virulence factors such as siderophores, phenazines and quorum sensing molecules that were previously shown to affect *C. albicans* growth (12). To test if known secreted factors contribute to the synergy seen with FLC and if they are transferable in conditioned media, we tested the activities of supernatants from WT *P. aeruginosa*, a double siderophore mutant and a phenazine mutant. Addition of the supernatant from *P. aeruginosa* did not affect *C. albicans* growth, whereas *P. aeruginosa* supernatant in combination with FLC completely blocked *C. albicans* trailing growth (Fig. 3A & Fig. S7). This is intermediate between FLC treatment alone, which results in trailing growth, and live *P. aeruginosa*, which synergizes to cause fungal death (Fig. 3A & Fig. S7). Conditioned media from *C. albicans* had no effect in combination with FLC, suggesting that the activity of *P. aeruginosa* supernatant is not due to a lack of nutrients, but rather from the activity of *P. aeruginosa*-secreted factors. Interestingly, the supernatant was not nearly as effective as live *P. aeruginosa* in synergizing with FLC. Supernatant from the double siderophore mutant and phenazine mutant strains performed indistinguishably from wildtype supernatant, inhibiting *C. albicans* trailing growth beyond the starting inoculum (Fig. 3B-D). Surprisingly, iron supplementation does not rescue *C. albicans* from the combination of FLC and *P. aeruginosa* conditioned media (Fig. S8). These results suggest that the full effects of *P. aeruginosa*-FLC synergy require live bacteria, while some activity can be transferred in conditioned media.

**Figure 3:**
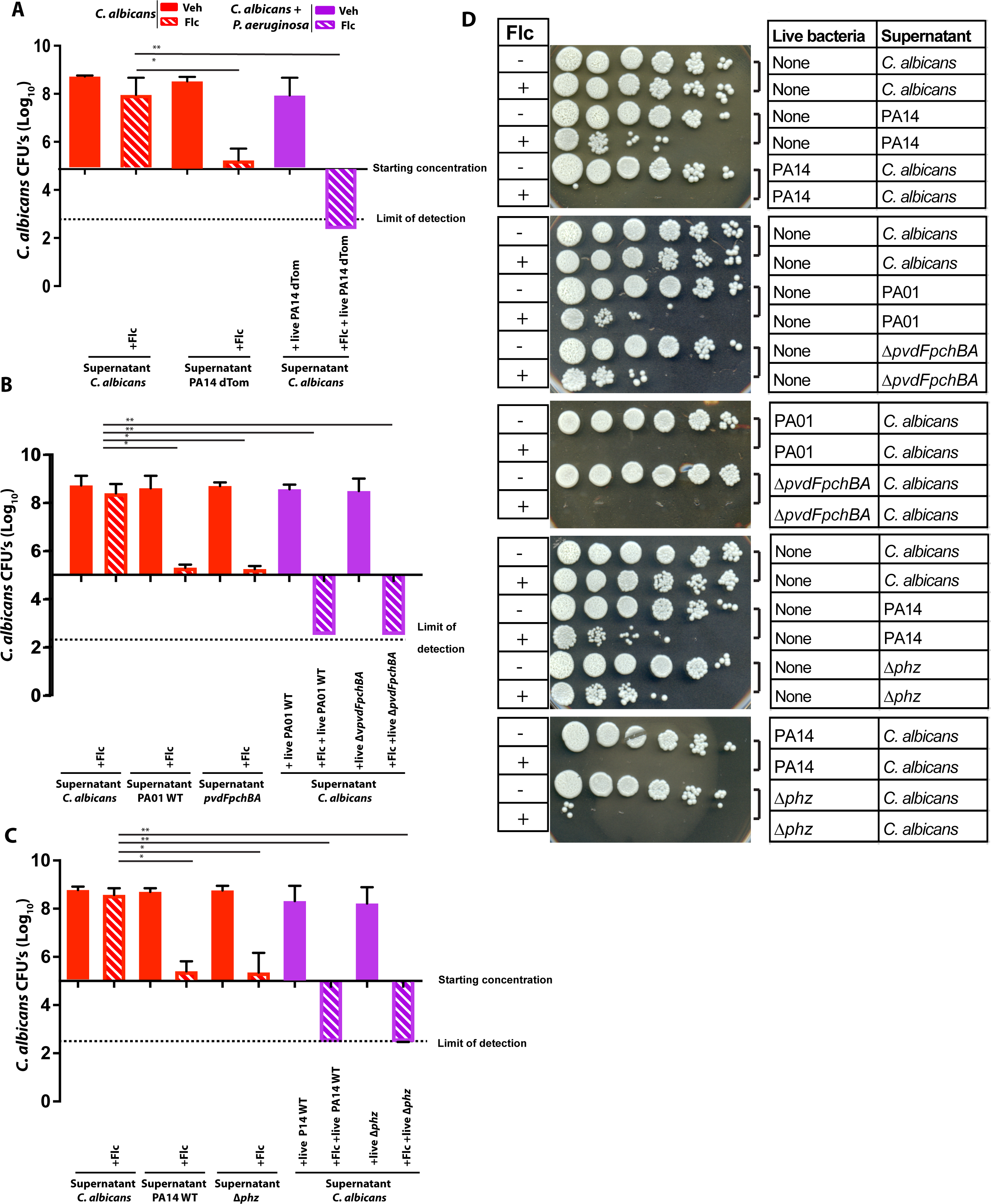
*P. aeruginosa* supernatant mild synergy with FLC compared to live *Pseudomonas*. *P. aeruginosa* and *C. albicans* were grown overnight in YPD media at 30°C. Overnight cultures supernatants were sterile filtered and added to 4 x 10^5^ *C. albicans* in YPD liquid media along with 12.5 µg/ml of FLC. After 48 hours of incubation at 30°C, cultures were 10-fold diluted and spotted onto YPD plates with antibiotics to count CFU. **A)** Supernatant from PA14-dTom strain, **B)** Supernatant from PA01 WT and Δ*pvdFpchBA*, **C)** Supernatant from PA14 WT and Δ*phz*, **D)** Representative images of YPD plates showing the growth of *C. albicans* after 24 hr of incubation. Data from 3 independent experiments. (p >0.05 NS; < 0.05 *; <0.01 **; <0.001 ***; <0.0001 ****)

### Iron homeostasis plays a limited role in regulation of *P. aeruginosa*-mediated synergy with FLC

To test the contribution of iron homeostasis to FLC-*P. aeruginosa* synergy during infection, we again turned to the zebrafish swimbladder model. We treated co-infections with FLC, supplemented with different levels of iron, and monitored both fish survival and fungal burden. Remarkably, iron supplementation reduced the protective effects of FLC against co-infection-induced mortality in a dose-dependent manner (Fig. 4A). Imaging revealed an increase in filamentous fungi that is usually associated with virulence but would be undercounted by homogenization and plating (23, 50, 56). To quantify this type of fungal overgrowth, we used double-blind scoring of individual fish for their level of hyphal growth, classifying fish into four categories (Fig. 4B). This semi-quantitative scoring revealed a mild but significant enhancement of fungal filamentous growth upon iron supplementation (Fig. 4C). This is also seen clearly in representative images selected from fish with median scores (Fig. 4D). Consistent with both the mild effect of siderophore deletion on FLC synergy *in vitro* and the intermediate effect of iron supplementation *in vivo*, the siderophore double mutant *P. aeruginosa* was not hypovirulent and did not limit the effectiveness of FLC during co-infection (Fig. 4E). Taken together with our *in vitro* findings, these *in vivo* infection results argue that iron supplementation has a limited ability to reverse *P. aeruginosa*-FLC synergy.

**Figure 4:**
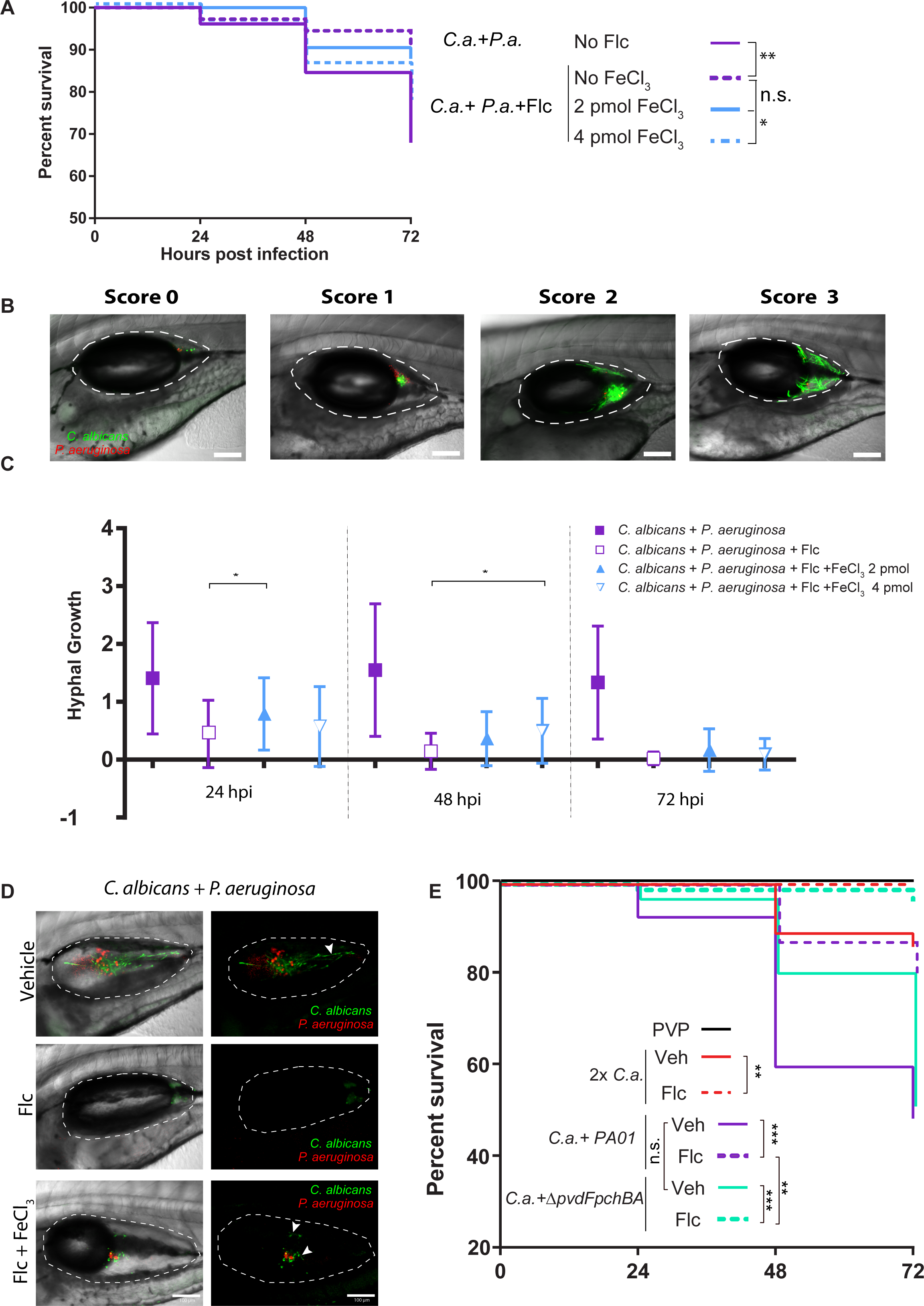
Iron homeostasis contributes to *P. aeruginosa*-mediated synergy with FLC. **A)** FeCl_3_ supplementation partially reverses *P. aeruginosa*-FLC synergy *in vivo*. Zebrafish injected with indicated microbes in the swimbladder with or without the indicated amounts of FeCl_3_.(2 or 4 pmol) Data pooled from 4 independent experiments. **B)** Hyphal growth during infection was scored using double-blind methodology. Representative images of each score: 0-no hyphal growth; 1-<10% coverage of swimbladder; 2-10-50% coverage of swimbladder; 3-> 50% coverage of swimbladder. **C)** FeCl_3_ supplementation is associated with stronger hyphal growth *in vivo*. Data shown are the medians with interquartile ranges from three experiments. **D)** Representative images of scored hyphal growth in the swimbladder at 24 hpi. Shown are median fish from each cohort. **E)** FLC treatment has no loss of effectiveness in co-infections with *P. aeruginosa* siderophore mutant. Scale bars = 100 μm. (p >0.05 NS; < 0.05 *; <0.01 **; <0.001 ***; <0.0001 ****)

## Discussion

In this study, we found that fungal-bacterial interactions can drive an unexpected enhancement in antifungal susceptibility during treatment of infection. Specifically, *P. aeruginosa* potentiates the fungicidal activity of the normally fungistatic drug fluconazole against *C. albicans*. We used a transparent mucosal infection model to mimic the clinical co-infections seen in cystic fibrosis and leveraged its simplicity and amenability to intravital imaging to probe the four-part interplay of two microbial species, drug therapy and host responses. These findings are clinically relevant for several reasons. First, these two microbes are frequently found together commensally and during infection, especially in cystic fibrosis. Second, there is a scarcity of effective anti-fungals and the action of the most orally bioavailable drug is limited by tolerance, which is associated with treatment failure. Third, our results implicate iron in infection and therapy in a new way beyond strictly as a micronutrient subjected to sequestration by the host and pathogen. The ability of *P. aeruginosa* to modify fungal drug susceptibility *in vitro* and *in vivo* adds a new dimension to the complexities of polymicrobial infection and raises importantquestions about the utilization of antifungal drugs during co-infection.

The fungicidal effect of FLC during co-infection suggests that *Pseudomonas* blocks *C. albicans* tolerance to FLC, leading to death rather than persistence or slow growth during treatment. Recent work suggests that drug tolerance should be considered alongside the traditional minimum inhibitory concentration (MIC) as an indicator for clinical response—and may be even more important than MIC (33). Determination of clinically-relevant drug resistance profiles in fungi is fraught with challenges and current *in vitro* testing protocols do not robustly match empirical clinical efficacy (32, 33). The disconnect between *in vitro* testing and clinical success may be due to biotic and/or abiotic factors in the host environment or may be due to a focus on the wrong metric for resistance. Microbe-microbe cross-talk alters antibacterial sensitivity *in vivo* (29, 57) and may be especially relevant in chronic co-infection of the immunocompromised host (13, 31). Thus, understanding how *P. aeruginosa* can reduce antifungal drug tolerance during treatment of infection has potentially important implications for both diagnosis and treatment.

Manipulation of iron homeostasis by *P. aeruginosa* is clearly one mechanism for altering FLC efficacy against *C. albicans*. This activity may be different from the iron piracy used by *P. aeruginosa* against other fungi, where *in vitro* antagonism is largely transferable with soluble factors such as siderophores (17, 58, 59). We tested other potentially contributing bacterial factors, including phenazines and quorum sensing, and fungal factors, including filamentous growth, but only iron supplementation significantly modulated the live *P. aeruginosa*-FLC synergy. Iron starvation is known to change FLC into a fungicidal drug, perhaps by regulating membrane fluidity, limiting mitochondrial function and/or blocking calcineurin-mediated stress responses (20, 33, 38, 55, 60). However, iron supplementation only partially reverses the effect of *P. aeruginosa* on FLC fungicidal activity both *in vitro* and during infection, and deletion of both major siderophores has minimal effects *in vitro* and no effect *in vivo*. Thus, while it is clear that *P. aeruginosa* has synergy with FLC against *C. albicans in vitro* and during co-infection, iron homeostasis is only one piece of the puzzle.

Iron is an essential micronutrient for both *P. aeruginosa* and *C. albicans* that each microbe acquires by multiple pathways during infection (15, 61, 62). Iron is important for *C. albicans* virulence in disseminated murine candidiasis and for epithelial invasion *in vitro* (63). Further, some iron chelators can work alone or in conjunction with FLC in murine models of OPC, VVC and disseminated mucormycosis (14, 39, 64–66). Conversely, iron supplementation can enhance virulence in both our zebrafish model and in a murine GI disease model (14, 25). Nonetheless, clinical studies of iron chelation against fungal infection are inconclusive and suggest it may negatively impact health (67–69). Thus, the prospect of using iron chelation to increase drug effectiveness during treatment of human patients holds both risks and potential benefits.

*C. albicans* can have both positive and negative interactions with diverse bacteria, depending on the context (12, 22, 70). In co-infection, *C. albicans* and *P. aeruginosa* are synergistically virulent in both a burn model and the zebrafish mucosal model (23, 26). However, these two microbes can exhibit antagonism *in vitro* and *in vivo* (12, 71). Furthermore, interactions of *C. albicans* with *S. epidermidis* and *S. aureus in vivo* are synergistic in terms of virulence but have shown antibiotic antagonism rather than synergy (72, 73). Given these disparate results, it remains to be tested whether the *P. aeruginosa*-FLC synergy is broadly relevant for vertebrate co-infections with *Candida*. It will be crucial to determine which mechanisms, beyond iron homeostasis, regulate *P. aeruginosa*-FLC synergy *in vitro* and then test those mechanisms in zebrafish and additional infection models such as the mouse cornea (74–76).

The ability of *P. aeruginosa* to synergize with FLC against *Candida* is only partly transferable in conditioned media, suggesting that there are multiple bacterial contributors to this ability. Interestingly, the effect of conditioned media does not depend on iron-chelating siderophores, and iron supplementation does not reverse the effect of conditioned media. Taken together, these data suggest that siderophores and iron starvation are only effective in synergizing with FLC when live bacteria are present to scavenge the iron-replete siderophores from the media highlighting the multifactorial nature of *P. aeruginosa* antagonism towards *Candida*.

Co-infections of *P. aeruginosa* and *C. albicans* are infrequent, except in the context of chronically infected cystic fibrosis (CF) patients and those on a ventilator (10, 77, 78). In CF, co-isolation of both *C. albicans* and *P. aeruginosa* is associated with worse outcomes, and the risks of other co-infections and acquisition of drug resistance are higher (13, 31). Interestingly, iron levels have been shown to be increased in cystic fibrosis airways and have been implicated in facilitating *P. aeruginosa* infections (79). Work in a zebrafish model of CF suggests that *P. aeruginosa* is similarly more virulent in the absence of CFTR activity in this vertebrate, suggesting that this model may be informative in translation to human disease (80).

In summary, FLC and *P. aeruginosa* have a synergistic interaction against *C. albicans* that results in enhanced clearance of *C. albicans*. This increased efficacy of FLC is dependent, in part, on iron sequestration caused by *P. aeruginosa*. We do not yet know if other *P. aeruginosa* clinical isolates show similar effect or if the synergy also occurs against other *Candida* species that are intrinsically more resistant to azoles, although a similar phenomenon occurs with *C. glabrata*. Nonetheless, our work demonstrates that polymicrobial interactions can profoundly shift antifungal sensitivity of *C. albicans*. On a more general level, our results also suggest that the biotic and abiotic growth environment can influence the efficacy of antifungal drugs, pointing the way towards new strategies for developing drugs to eradicate recalcitrant infections.

## Materials and Methods

### Ethics Statement and Zebrafish care and maintenance

Adult zebrafish used for breeding embryos were housed in recirculating systems (Aquatic Habitats, Apopka, FL) at the University of Maine Zebrafish Facility. All zebrafish care protocols and experiments were performed in accordance with National Research Council guidelines (81) under Institutional Animal Care and Use Committee (IACUC) protocols A2015-11-03 and A2018-10-01. Larvae were reared at a density of 150/dish in 150-mm petri dishes containing 150 ml of E3 (5 mM sodium chloride, 0.174 mM potassium chloride, 0.33 mM calcium chloride, 0.332 mM magnesium sulfate, 2 mM HEPES in Nanopure water, pH 7) supplemented with 0.02 mg/ml of 1-phenyl-2-thiourea (PTU) (Sigma-Aldrich, St. Louis, MO) to prevent pigmentation, as well as 0.3 mg/liter methylene blue (VWR, Radnor, PA) for the first 24 h to prevent microbial growth. Larvae were manually dechorionated at 24 h postfertilization, transferred into media containing E3 and PTU, and incubated at 33°C over the course of experiments. This temperature was chosen as the highest safe temperature for zebrafish health to approximate human body temperature and is regularly used for experiments with temperature-sensitive alleles. Experiments were conducted using wild-type (AB) zebrafish.

### Strains and growth conditions

The strains used in this study are listed in Table 1. Most experiments were conducted with the *C. albicans* reference strain SC5314 and either PA14 or PA01 *P. aeruginosa* strains. All experiments with mutant bacteria or fungi were conducted with matched controls from the source laboratory (see Table S1 for panel-by-panel description). *C. albicans* and *P. aeruginosa* strains were routinely refreshed from frozen stocks at -80°C and maintained on YPD (1% Bacto yeast extract, 2% Bacto peptone, 2% dextrose, 2% Bacto agar) plates or *Pseudomonas* isolation agar (Sigma-Aldrich) for *in vitro* experiments and LB agar (10 g/liter Bacto tryptone, 5 g/liter Bacto yeast extract, 10 g/liter sodium chloride, 1.2% agar; BD, San Jose, CA) supplemented with 750 μg/ml ampicillin (EMD Millipore, Billerica, MA)for injection. Liquid cultures were grown overnight in YPD or LB media in a rotator wheel at 30°C. Prior to experiments, cultures were washed with phosphate-buffered saline (PBS) and optical density OD 600 was measured.

**Table 1.** Fungal and bacterial strains used

| Strain Name | Description and Genotype | Reference |
| --- | --- | --- |
| ***Candida albicans* and *Candida glabrata*** |  |  |
| SC5314-Neon | Wildtype clinical isolate, pENO1-NEON-NAT | (82) |
| Caf2-FR | SC5314 background; Δura3::imm434/URA3 pENO1-iRFP-NAT | (23) |
| SN250 | *his1*Δ/*his1*Δ, *leu2*Δ::*C.dubliniensis* HIS1/*leu2*Δ::*C.maltosa* LEU2, *arg4*Δ /*arg4*Δ, URA3/*ura3*Δ::imm434, *IRO1*/*iro1*Δ::imm434 | (83) |
| *sef1* $\Delta/\Delta$ | *his1*Δ/*his1*Δ, *sef1*Δ::*C.dubliniensis* HIS1/*sef1*Δ::*C.maltosa* LEU2, *arg4*Δ /*arg4*Δ, URA3/*ura3*Δ::imm434, *IRO1*/*iro1*Δ::imm434 | (83) |
| *sfu1* $\Delta/\Delta$ | *his1*Δ/*his1*Δ, *sfu1*Δ::*C.dubliniensis* HIS1/*sfu1*Δ::*C.maltosa* LEU2, *arg4*Δ /*arg4*Δ, URA3/*ura3*Δ::imm434, *IRO1*/*iro1*Δ::imm434 | (83) |
| NCO-788 | Clinical isolate | (84) (Clancy and Shields, U. Pittsburgh) |
| NC1 *C. glabrata* | Clinical isolate | (84) (Clancy and Shields, U. Pittsburgh) |
| NC999 *C. glabrata* | Clinical isolate | (84) (Clancy and Shields, U. Pittsburgh) |
| CG-4720 *C. glabrata* | Clinical isolate | (85) |
| B13-TWO7229#2 | Clinical isolate #2 in series from patient. | (84) (86) |
| B14-TWO7230#3 | Clinical isolate #3 in series from patient | (84) (86) |
| B15-TWO7241#16 | Clinical isolate #16 in series from patient | (84) (86) |
| B16-TWO7243#17 | Clinical isolate #17 in series from patient | (84) (86) |
| ***Pseudomonas aeruginosa*** |  |  |
| PA14 dTom | PA14 carrying plasmid encoding dTomato | (80) |
| PA14 $\Delta lasR$ | In-frame deletion of *lasR* | (87) |
| PA01 WT | Wild type clinical isolate | (88) |
| PAO6382 | PA01 $\Delta pvdF$ | (89) |
| PAO6297 | PA01 $\Delta pchBA$ | (90) |
| PAO6383 | PA01 $\Delta pvdF\Delta pchBA$ | (90) |
| PA14 WT | Wildtype clinical isolate | (91) |
| PA14 $\Delta phz$ | In-frame deletion of *phzA1-G1* and *phzA2-G2* operons, phenazine negative | (91) |
| PA14 *phzM::TnM* | TnM mutant, 5-MPCA negative | (91) |
| PA14 *phzS::TnM* | TnM mutant, PYO negative | (91) |

### In vitro C. albicans and P. aeruginosa co-culture

*P. aeruginosa* and *C. albicans* were individually grown overnight. *P. aeruginosa* was grown in GGP media (3% glycerol, 1% proteose peptone, 2.9 mM K_2_HPO_4_, 2 mM MgSO_4_●7H_2_0) or LB media (10 g/liter Bacto tryptone, 5 g/liter Bacto yeast extract, 10 g/liter sodium chloride; BD, San Jose, CA) at 30°C. *C. albicans* was grown in YPD media at 30°C. *P. aeruginosa* and *C. albicans* cultures were combined in a 1:1 ratio with both organisms at a final concentration of 2 x 10^5^/ml. The *P. aeruginosa/C. albicans* coculture was grown at 30°C for 48 hours in YPD on a rotating wheel. The standard concentration of fluconazole used was 12.5 µg/ml (Sigma Aldrich). The spot test was performed by spotting 3 μl from each dilution using a multichannel pipette, plated on YPD agar supplemented with penicillin (250 U/ml)-streptomycin (250 μg/ml) (Lonza), 30 μg/ml gentamicin sulfate (BioWhittaker, Lonza), and 3 μg/ml vancomycin hydrochloride (Amresco, Solon, OH) and on *Pseudomonas* isolation agar (Sigma-Aldrich) for *C. albicans* and *P. aeruginosa* selection respectively. Plates were incubated for 24 hours at 37 °C. For collecting supernatants, *P. aeruginosa* overnight cultures in LB or GGP were centrifuged at 21,000 x g for 2 minutes and supernatant was filtered using an Acrodisc 0.2 µm syringe filter (PALL corporation). Filtered supernatant was added to 4 x 10^5^ *C. albicans* in YPD liquid media. 48 hours post incubation, cultures were 10-fold diluted and spot tests were performed as described above.

### Swimbladder infections via microinjection

At 4 days postfertilization, zebrafish larvae were anesthetized in Tris-buffered tricaine methane sulfonate (160 μg/ml; Tricaine; Western Chemicals, Inc., Ferndale, WA) and selected for swimbladder inflation. Fish were microinjected as previously described (48). Fungal and bacterial cells were resuspended in 5% polyvinylpyrrolione (PVP; Sigma-Aldrich) and fish were injected with 4 nl of PVP control, *C. albicans* at 5 x 10^7^ CFU/ml, or a *C. albicans-P. aeruginosa* mixture at 2.5 x 10^7^ CFU/ml for each. The *C. albicans-P. aeruginosa* coculture was prepared by combining equal volumes of *C. albicans* at 5 x 10^7^ CFU/ml and *P. aeruginosa* at 5 x 10^7^ CFU/ml prior to injection. As indicated, FeCl_3_ (Sigma-Aldrich) was added to the injection solution with *C. albicans* and/or *P. aeruginosa* to a final concentration of 0.5 mM or 1 mM, for a final amount of 2 or 4 pmol per 4 nl dose.

Within 1 hr of injection, larvae were placed in individual wells of a 96-well glass-bottom imaging dish (Greiner Bio-One, Monroe, NC) and screened for an inoculum of 50 to 100 yeast cells for mono-infection, and 25 to 50 yeast cells for co-infection, using a Zeiss AxioVision VivaTome microscope. For mortality experiments, fish were kept at 33°C in E3 containing PTU with or without fluconazole at 100 μg/ml. Fish were held for 3 days post injection and monitored daily for survival.

### Confocal laser scanning fluorescence microscopy

At 24, 48 and 72 h post-injection, larvae were anesthetized in Tricaine and immobilized in 0.4 % low-melting-point agarose (Lonza, Switzerland) in E3 containing Tricaine in a 96-well glass-bottom imaging dish (Greiner Bio-One, Monroe, NC). Confocal images were acquired using an Olympus IX-81 inverted microscope with an FV-1000 laser scanning confocal system (Olympus, Waltham, MA). The EGFP, dTomato, and Far-Red fluorescent proteins were detected by laser/optical filters with a 20x (NA, 0.7) for excitation/emission at 488 nm/505 to 525 nm, 543 nm/560 to 620 nm, and 635 nm/655 to 755 nm, respectively. Z-stacks of 15 to 25 slices, with an interslice interval between 7 and 13 μm, were collected and processed using FluoView (Olympus, Waltham, MA)

### Image analysis

The percentage of the swimbladder covered by *Candida* at 24, 48 and 72 hpi was determined using Fiji software (ImageJ) applied to maximum-projection images from stacks of 15 to 25 z-slices. Images were taken with identical acquisition settings to ensure comparability. The swimbladder area was delineated, and the percent coverage of *Candida* fluorescence above a set threshold (corresponding to background fluorescence) was calculated. Images covered the swimbladder from midline to skin in 5-μm z-slices. The number of slices per image ranged from 15 to 25, depending on the size of the fish.

### Hyphal growth scoring from confocal images

Zebrafish infected with *C. albicans* SC5314-Neon and PA14-dTomato with or without Fluconazole. FeCl_3_ (2 or 4 pmol) was co-injected into swimbladder along with *C. albicans and P. aeruginosa.* Fish were imaged at 24, 48 and 72 hpi and images were processed as described above. Hyphal growth of *C. albicans* in the swimbladder was scored blindly as follows; Score 0 for no hyphal growth. Score 1 for <10% hyphal growth. Score 2 between 10 and 50% hyphal growth. Score 3 > 50% hyphal growth.

### CFU assessments

For CFU quantification, 5 randomly selected infected larvae were pooled and homogenized at 24, 48 and 72 hpi in 500 μl of 1X PBS. For plating, 50 μl or 100 μl of homogenate from groups was plated on both YPD agar supplemented with 250 U/ml ,250 μg/ml penicillin-streptomycin (Lonza), 30 μg/ml gentamicin sulfate (BioWhittaker, Lonza), and 3 μg/ml vancomycin hydrochloride (Amresco, Solon, OH) and on *Pseudomonas* isolation agar (Sigma-Aldrich) for *C. albicans* and *P. aeruginosa* selection, respectively. Plates were incubated overnight at 37°C, colonies were counted the following day, and CFU/fish was calculated.

### Statistical analyses

Statistical analyses were conducted using GraphPad Prism 7 software (GraphPad Software, Inc., La Jolla, CA). Data was analyzed for normality and appropriate parametric or nonparametric tests were performed, means or medians are shown, respectively. All significant differences are indicated in the figures, with *, **, ***, and **** indicating P values of <0.05, <0.01, <0.001, and <0.0001, respectively. Kaplan-Meier survival curves were subjected to a log rank (Mantel-Cox) test, and Bonferroni correction was then used to determine statistical differences between pairs of treatments. Monte-Carlo simulation was used to analyze ratios in Fig 1E. Mann-Whitney test was used to analyze experiments in figure panels 1D, 1F, 2A, 4C. Unpaired T-test was used for Fig. 1F. Two-way ANOVA was used for Fig. 2B. For Fig. 3A-C, significance was established by identifying non-overlapping 95% confidence intervals.

## Acknowledgements

We would like to thank Dr. Damian Krysan (U. Iowa) for providing *Candida* clinical isolates, Dr. Natalia Kirienko (Rice University), Dr. Deborah Hogan (Dartmouth University) and Dr. Andrew Koh (UT Southwestern) for providing *P. aeruginosa* mutants. We would like to thank Dr. Robert Shanks (U. Pittsburgh) for consultation and providing initial *P. aeruginosa* supernatants. We would like to thank Mark Nilan for outstanding zebrafish husbandry, members of the Wheeler Laboratory and Dr. Gerry Fink for comments on the manuscript, and several undergraduates for their useful contributions (Roxane Baudouin, Orlane Mombled, Jennifer Quezada-Loja, Jessica Hayden, Maria Vina Lopez, and Nikhil Vaidya). R.T.W. is a Burroughs Wellcome Fund investigator in the pathogenesis of infectious disease, S.H. is Chase fellow at UMaine, and this work was funded by NIH grant R15AI133415 and by the USDA National Institute of Food and Agriculture, Hatch project number ME0-21821, through the Maine Agricultural and Forest Experiment Station.

## Supplemental Information

### Supplemental Methods

#### Statistical analyses

Statistical analyses were conducted using GraphPad Prism 7 software (GraphPad Software, Inc., La Jolla, CA). Data was analyzed for normality and appropriate parametric or nonparametric tests were done; means or medians are shown, respectively. All significant differences are indicated in the figures, with *, **, ***, and **** indicating P values of <0.05, <0.01, <0.001, and <0.0001, respectively. Kaplan-Meier survival curves were subjected to a log rank (Mantel-Cox) test, and Bonferroni correction was then used to determine statistical differences between pairs of treatments. One-way ANOVA was used to analyze Fig. S2, S3 and S5. Two-way ANOVA was used for Fig. S6. For Fig. S7 and Fig. S8, significance was established by identifying non-overlapping 95% confidence intervals.

### Supplemental Figure Legends

**Fig. S1. *P. aeruginosa* enhances FLC activity against *C. albicans* fluconazole-resistant strains and *C. glabrata* FLC hyper resistant strains.**

MIC_50_ was tested separately and supra-MIC concentrations of FLC were used (per measurements relevant for our assays; Table S1). *Candida* clinical isolates and *P. aeruginosa* PA14 were co-cultured in YPD. FLC was added at concentrations above the MIC_50_. Co-cultures were incubated at 30° C for 48 hr, then 10x dilutions of the co-cultures were spotted on YPD agar containing antibiotics. Pictures were taken after 24 hr of incubation at 37°C. Representative results of three independent experiments are shown.

**Supplemental Table 1.** Relevant measured MIC_50_ for clinical strains

| Clinical isolate strain | FLC MIC<sub>50</sub> as measured here |
| --- | --- |
| SN250 (WT) | 12.5 µg/ml |
| NCO-788 | 200 µg/ml |
| NC1 *C. glabrata* | 50 µg/ml |
| NC999 *C. glabrata* | 400 µg/ml |
| CG-4720 *C. glabrata* | 200 µg/ml |
| B13 TWO7229#2 | 50 µg/ml |
| B14 TWO7230#3 | 50 µg/ml |
| B15 TWO7241#16 | 100 µg/ml |
| B16 TWO7243#17 | 400 µg/ml |

**Fig. S2. Heat killed *Pseudomonas aeruginosa* does not synergize with FLC *in vitro***.

*P. aeruginosa* was heat killed by incubating at 100*°* C for 20 min. Heat killed *P. aeruginosa* was adjusted to 2 x 10^7^/ml in YPD and added to 2 x 10^5^/ml of *C. albicans*. Co-cultures were incubated at 30° C for 48 hr. 3 μl of serial 10-fold dilutions of co-cultures were plated on YPD agar plate containing antibiotics or on PIA plates selective for *P. aeruginosa*. Data shown are the medians with ranges for three experiments.

**Fig. S3. *P. aeruginosa* also synergizes with FLC at 37°C and in YPD + FBS.** (A) Fluconazole (FLC) treatment of *C. albicans* and *P. aeruginosa* co-culture shows a fungicidal effect in YPD at 37*°* C. *P. aeruginosa* and *C. albicans* were inoculated at 2x10^5^/ml in YPD, FLC was added at 12.5 μg/ml. After incubation of liquid co-cultures at 37*°*C for 48 hr, 3 μl of serial 10-fold dilutions of co-cultures were plated on YPD containing antibiotics. Plates were grown at 37*°* C for 24 hr. Data shown are the medians with ranges for three experiments. (B-C) Fluconazole (FLC) treatment of *C. albicans* and *P. aeruginosa* shows a fungicidal effect after co-culture in YPD + 2% FBS at 37*°* C. *P. aeruginosa* and *C. albicans* were inoculated at 2x10^5^/ml in YPD + 2% FBS. FLC was added at 12.5 μg/ml and liquid cultures incubated at 37*°*C for 48 hr. (B) 3 μl of serial 10-fold dilutions of co-cultures were plated on YPD containing antibiotics. Plates were grown at 37*°* C for 24 hr before counting. Data shown are the medians with ranges from three experiments. (C) Representative images showing *C. albicans* morphology as yeast in YPD or hyphal growth in YPD + 2% FBS. After 48 hr of co-culture, 5 μl of co-culture was imaged using Zeiss Vivatome microscope. White arrowheads show hyphal growth in YPD +2 % FBS. Black arrowheads show swollen cells associated with trailing growth in FLC. Arrows note dead *C. albicans* cells in both YPD and YPD + 2% FBS conditions in co-cultures with FLC.

**Fig. S4. FLC does not affect larval zebrafish survival**. Zebrafish larvae (4 dpf) were anesthetized in Tris-buffered tricaine methane sulfonate and selected for swimbladder inflation. Fish were microinjected with a 4 nl of PVP control. Fish were kept at 33°C in E3 containing PTU with or without FLC at 100 μg/ml. Fish were held for 3 days post injection and monitored daily for survival.

**Fig. S5. FLC does not affect *P. aeruginosa* growth *in vitro***. Addition of FLC to *P. aeruginosa* alone and in co-culture with *C. albicans* does not affect *P. aeruginosa* growth *in vitro*. *P. aeruginosa* and *C. albicans* were inoculated at 2x10^5^/ml in YPD, FLC was added at 12.5 μg/ml . Cultures were grown at 30° C for 48 hr. 3 μl of serial 10-fold dilutions of co-cultures were plated on *Pseudomonas* isolation agar (PIA). Graph showing log_10_ CFU/ml for *P. aeruginosa* viability. Data shown are the median with ranges from 3 experiments.

**Fig S6: Phenazines and quorum sensing molecule LasR do not contribute to the synergistic activity with FLC. A)** Co-culture of *C. albicans* with *P. aeruginosa* WT or Δ*lasR* mutant PA14 Δ*lasR* mutant is synergistic with FLC. Bar graph represents *C. albicans* growth in log_10_ CFU/ml. Data is representative of 3 independent experiments. **B)** Co-culture of *C. albicans* with *P. aeruginosa* WT or phenazine deficient strains: PA14 Δ*phz,* PA14 *phzM::TnM ,*PA14 *phzS::TnM* in the presence or absence of FLC (12.5 μg/ml). Bar graph represents *C. albicans* growth in log_10_ CFU/ml. Data is representative of 3 independent experiments. Data shown are the median with ranges.

**Fig. S7. *P. aeruginosa* supernatant affect *C. albicans* trailing growth in the presence of FLC**

Addition of *P. aeruginosa* supernatant is not as effective as live *P. aeruginosa* in inhibiting *C. albicans* growth, but supernatant from both PA14 WT and PA14 Δ*phz* causes inhibition of trailing growth past the starting inoculum. Graph showing *C. albicans* viability after 24 hr and 48 hr of growth in the presence of FLC, with *C. albicans* or *P. aeruginosa* supernatant, with orwithout live *P. aeruginosa*. Data show medians and 95% confidence intervals from 3 experiments, with comparable data from other experiments with siderophore mutants.

**Fig. S8. FeCl_3_ supplementation does not block fungicidal synergy of *P. aeruginosa* combined with FLC.**

FeCl_3_ supplementation reverses fungicidal activity of live *P. aeruginosa* + FLC, but not when added to *P. aeruginosa* supernatant + FLC. Supernatant from overnight cultures of either *C. albicans* or *P. aeruginosa* grown in YPD was filter-sterilized, and added at 1:1 dilution to YPD with *C. albicans* at 2x10^5^/ml. FLC was added at 12.5 μg/ml and different concentrations of FeCl_3_ used range from 1 mM to 0.125 mM. Data are representative of 3 independent experiments. Graphs show medians with 95% confidence intervals.

